# *TP53* mutations and drug sensitivity in acute myeloid leukaemia cells with acquired MDM2 inhibitor resistance

**DOI:** 10.1101/404475

**Authors:** Martin Michaelis, Constanze Schneider, Florian Rothweiler, Tamara Rothenburger, Marco Mernberger, Andrea Nist, Andreas von Deimling, Daniel Speidel, Thorsten Stiewe, Jindrich Cinatl

## Abstract

**Background:** MDM2 inhibitors are under investigation for the treatment of acute myeloid leukaemia (AML) patients in phase III clinical trials. To study resistance formation to MDM2 inhibitors in AML cells, we here established 45 sub-lines of the AML *TP53* wild-type cell lines MV4-11 (15 sub-lines), OCI-AML-2 (10 sub-lines), OCI-AML-3 (12 sub-lines), and SIG-M5 (8 sub-lines) with resistance to the MDM2 inhibitor nutlin-3.

Methods: Nutlin-3-resistant sub-lines were established by continuous exposure to stepwise increasing drug concentrations. The *TP53* status was determined by next generation sequencing, cell viability was measured by MTT assay, and p53 was depleted using lentiviral vectors encoding shRNA.

**Results:** All MV4-11 sub-lines harboured the same R248W mutation and all OCI-AML-2 sub-lines the same Y220C mutation, indicating the selection of pre-existing *TP53*-mutant subpopulations. In concordance, rare alleles harbouring the respective mutations could be detected in the parental MV4-11 and OCI-AML-2 cell lines. The OCI-AML-3 and SIG-M5 sub-lines were characterised by varying *TP53* mutations or wild type *TP53*, indicating the induction of *de novo TP53* mutations. Doxorubicin, etoposide, gemcitabine, cytarabine, and fludarabine resistance profiles revealed a noticeable heterogeneity among the sub-lines even of the same parental cell lines. Loss-of-p53 function was not generally associated with decreased sensitivity to cytotoxic drugs.

**Conclusion:** We introduce a substantial set of models of acquired MDM2 inhibitor resistance in AML. MDM2 inhibitors select, in dependence on the nature of a given AML cell population, pre-existing *TP53*-mutant subpopulations or induce *de novo TP53* mutations. Although loss-of-p53 function has been associated with chemoresistance in AML, nutlin-3-adapted sub-lines displayed in the majority of experiments similar or increased drug sensitivity compared to the respective parental cells. Hence, chemotherapy may remain an option for AML patients after MDM2 inhibitor therapy failure. Even sub-lines of the same parental cancer cell line displayed considerable heterogeneity in their response to other anti-cancer drugs, indicating the need for the detailed understanding and monitoring of the evolutionary processes in cancer cell populations in response to therapy as part of future individualised treatment protocols.

## Background

MDM2 inhibitors are under development as novel class of anti-cancer drugs for the treatment *TP53* wild-type cancer cells from different cancer entities including acute myeloid leukaemia (AML) [1]. *TP53* encodes p53, a major tumour suppressor protein. *MDM2* is a p53 target gene that encodes for MDM2, a major endogenous inhibitor of p53. MDM2 physically interacts with p53 and mediates its ubiquitination and proteasomal degradation. MDM2 inhibitors activate p53 signalling by interference with the MDM2/ p53 interaction [1-3].

Various MDM2 inhibitors have been shown to exert anti-cancer effects in pre-clinical models of AML, alone or in combination with other drugs [4-20]. Moreover, different MDM2 inhibitors are under investigation in clinical studies for their effects on AML [18,21-23], with idasanutlin currently being tested in phase II and III trials for the treatment of AML (NCT02670044, NCT02545283).

Drug-adapted cancer cell lines have been used to identify and investigate clinical resistance mechanisms [24-33]. The adaptation of cancer cell lines to MDM2 inhibitors indicated that the treatment of *TP53* wild-type cancer cells may be associated with the formation of *TP53* mutations as resistance mechanisms [3,34-39]. In concordance, treatment of liposarcoma patients harbouring *TP53* wild type cancer cells with the MDM2 inhibitor SAR405838 resulted in the emergence of *TP53* mutations [40].

The origin of MDM2 inhibitor-induced *TP53* mutations in *TP53* wild-type cell lines is not entirely clear. In dependence of the cell line model, MDM2 inhibitors may induce a range of different *de novo TP53* mutations in a given model or select small, pre-existing cell fractions that harbour *TP53* mutations [35,36,39,41].

To study acquired resistance formation to MDM2 inhibitors in AML cells, we here established and analysed a panel of sub-lines of the *TP53* wild-type AML cell lines MV4-11, OCI-AML-2, OCI-AML-3, and SIG-M5, with acquired resistance to the MDM2 inhibitor nutlin-3 [3,42]. In total, this included 45 nutlin-3-adapted sub-lines (15 MV4-11 sub-lines, 10 OCI-AML-2 sub-lines, 12 OCI-AML-3 sub-lines, 8 SIG-M5 sub-lines).

## Methods

### Cells

The AML cell lines MV4-11, OCI-AML-2, OCI-AML-3, and SIG-M5 were obtained from DSMZ (Braunschweig, Germany). The nutlin-3-resistant sub-lines were established by adaption to growth in the presence of increasing drug concentrations as previously described [35,36] and derived from the resistant cancer cell line (RCCL) collection [43].

All cells were propagated in IMDM supplemented with 10 % FBS, 100 IU/mL penicillin and 100 µg/mL streptomycin at 37°C. Cells were routinely tested for mycoplasma contamination and authenticated by short tandem repeat profiling.

p53-depleted SIG-M5 cells were established as described previously [44] using the Lentiviral Gene Ontology (LeGO) vector technology [45,46].

### Viability assay

Cell viability was tested by the 3-(4,5-dimethylthiazol-2-yl)-2,5-diphenyltetrazolium bromide (MTT) dye reduction assay after 120 h incubation modified as described previously [35,36]. 2×10^4^ cells suspended in 100 µL cell culture medium were plated per well in 96-well plates and incubated in the presence of various drug concentrations for 120 h. Then, 25µL of MTT solution (2 mg/mL (w/v) in PBS) were added per well, and the plates were incubated at 37°C for an additional 4h. After this, the cells were lysed using 100µL of a buffer containing 20% (w/v) sodium dodecylsulfate and 50% (v/v) N,N-dimethylformamide with the pH adjusted to 4.7 at 37°C for 4h. Absorbance was determined at 560 nm to 620 nm for each well using a 96-well multiscanner. After subtracting of the background absorption, the results are expressed as percentage viability relative to control cultures which received no drug.

Drug concentrations that inhibited cell viability by 50% (IC50) were determined using CalcuSyn (Biosoft, Cambride, UK).

### *TP53* next generation sequencing

The *TP53* status was determined by next generation sequencing as previously described [47]. All coding exonic and flanking intronic regions of the human TP53 gene were amplified from genomic DNA with Platinum^TM^ Taq DNA polymerase (Life Technologies) by multiplex PCR using two primer pools with 12 non-overlapping primer pairs each, yielding approximately 180 bp amplicons. Each sample was tagged with a unique 8-nucleotide barcode combination using twelve differently barcoded forward and eight differently barcoded reverse primer pools. Barcoded PCR products from up to 96 samples were pooled, purified and an indexed sequencing library was prepared using the NEBNext® ChIP-Seq Library Prep Master Mix Set for Illumina in combination with NEBNext® Multiplex Oligos for Illumina (New England Biolabs). The quality of sequencing libraries was verified on a Bioanalyzer DNA High Sensitivity chip (Agilent) and quantified by digital PCR. 2 x 250 bp paired-end sequencing was carried out on an Illumina MiSeq (Illumina) according to the manufacturer’s recommendations at a mean coverage of 300x.

Read pairs were demultiplexed according to the forward and reverse primers and subsequently aligned using the Burrows-Wheeler Aligner against the Homo sapiens Ensembl reference (rev. 79). Overlapping mate pairs were combined and trimmed to the amplified region. Coverage for each amplicon was calculated via SAMtools (v1.1) [48]. To identify putative mutations, variant calling was performed using SAMtools in combination with VarScan2 (v2.3.9) [49]. Initially, SAMtools was used to create pileups with a base quality filter of 15. Duplicates, orphan reads, unmapped and secondary reads were excluded. Subsequently, Varscan2 was applied to screen for SNPs and InDels separately, using a low-stringency setting with minimal variant frequency of 0.1, a minimum coverage of 20 and a minimum of 10 supporting reads per variant to account for cellular and clonal heterogeneity. Minimum average quality was set to 20 and a strand filter was applied to minimize miscalls due to poor sequencing quality or amplification bias. The resulting list of putative variants was compared against the IARC TP53 (R17) database to check for known p53 cancer mutations.

### Statistics

Results are expressed as mean ± S.D. of at least three experiments. Comparisons between two groups were performed using Student’s t-test. Three and more groups were compared by ANOVA followed by the Student-Newman-Keuls test. P values lower than 0.05 were considered to be significant.

## Results

### Nutlin-3 sensitivity/ resistance status of the nutlin-3-adapted AML sub-lines

To study acquired resistance formation to MDM2 inhibitors in AML cells, we established and analysed a panel of sub-lines of the *TP53* wild-type AML cell lines MV4-11, OCI-AML-2, OCI-AML-3, and SIG-M5, with acquired resistance to the MDM2 inhibitor nutlin-3. The parental cell lines MV4-11, OCI-AML-2, OCI-AML-3, and SIG-M5 displayed sensitivity to nutlin-3 in a range of 0.90 to 2.33µM (Suppl. Table 1). The nutlin-3 IC50 values in the nutlin-3-adapted sub-lines of MV4-11 (nutlin-3 IC50: 2.33µM) ranged from 13.3 to 22.6µM resulting in resistance factors (fold change nutlin-3 IC50 in nutlin-3-adapted MV4-11 sub-lines/ nutlin-3 IC50 in MV4-11) ranging between 5.7 (MV4-11^r^Nutlin^20µM^XII) and 9.7 (MV4-11^r^Nutlin^20µM^II) (Figure 1, Suppl. Table 1).

**Table 1.**
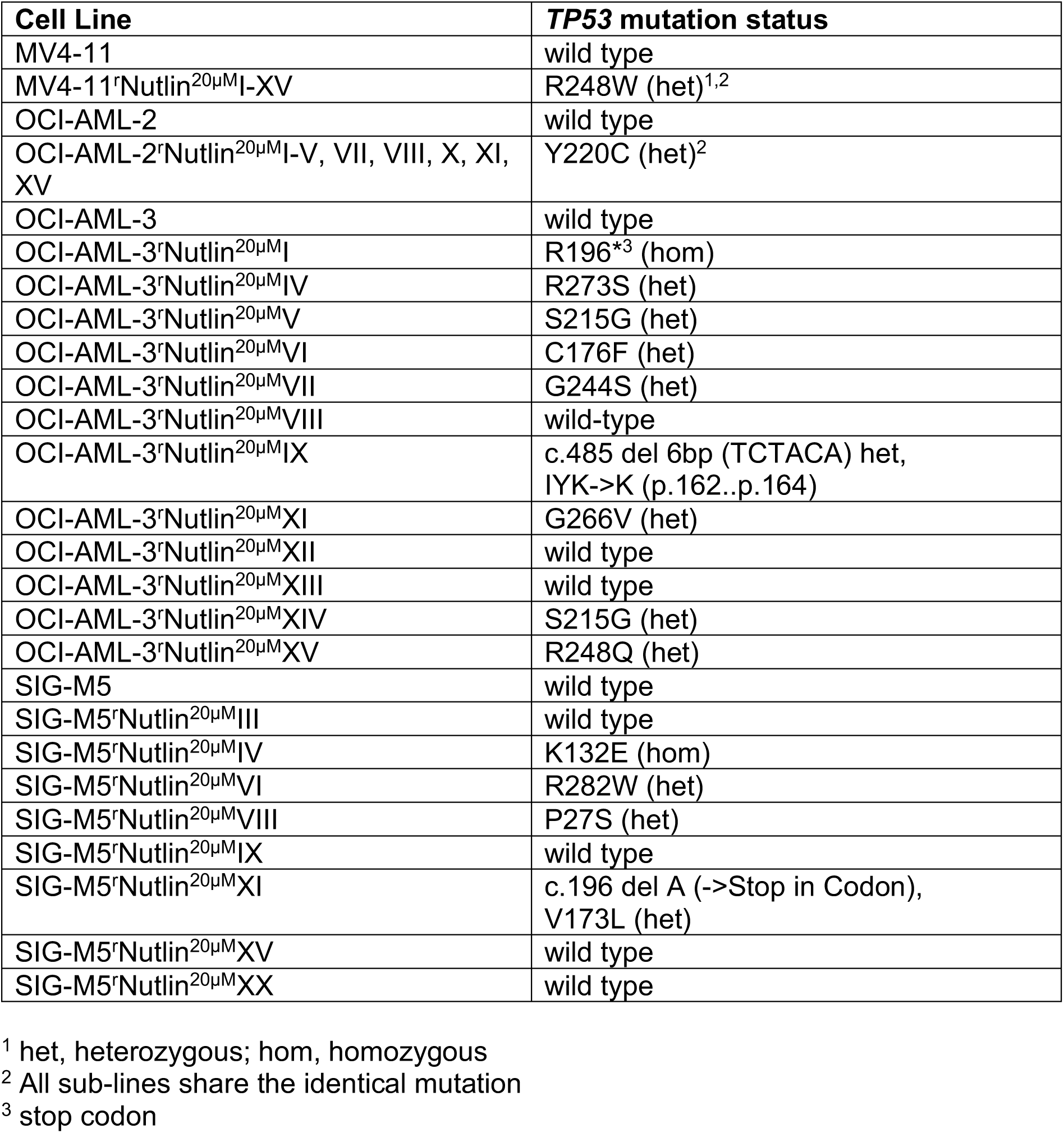
*TP53* mutation status of AML cell lines and their nutlin-3-adapted sub-lines.

| Cell Line | *TP53* mutation status |
| --- | --- |
| MV4-11 | wild type |
| MV4-11rNutlin<sup>20μM</sup>I-XV | R248W (het)<sup>1,2</sup> |
| OCI-AML-2 | wild type |
| OCI-AML-2rNutlin<sup>20μM</sup>I-V, VII, VIII, X, XI, XV | Y220C (het)<sup>2</sup> |
| OCI-AML-3 | wild type |
| OCI-AML-3rNutlin<sup>20μM</sup>I | R196*<sup>3</sup> (hom) |
| OCI-AML-3rNutlin<sup>20μM</sup>IV | R273S (het) |
| OCI-AML-3rNutlin<sup>20μM</sup>V | S215G (het) |
| OCI-AML-3rNutlin<sup>20μM</sup>VI | C176F (het) |
| OCI-AML-3rNutlin<sup>20μM</sup>VII | G244S (het) |
| OCI-AML-3rNutlin<sup>20μM</sup>VIII | wild-type |
| OCI-AML-3rNutlin<sup>20μM</sup>IX | c.485 del 6bp (TCTACA) het, IYK->K (p.162..p.164) |
| OCI-AML-3rNutlin<sup>20μM</sup>XI | G266V (het) |
| OCI-AML-3rNutlin<sup>20μM</sup>XII | wild type |
| OCI-AML-3rNutlin<sup>20μM</sup>XIII | wild type |
| OCI-AML-3rNutlin<sup>20μM</sup>XIV | S215G (het) |
| OCI-AML-3rNutlin<sup>20μM</sup>XV | R248Q (het) |
| SIG-M5 | wild type |
| SIG-M5rNutlin<sup>20μM</sup>III | wild type |
| SIG-M5rNutlin<sup>20μM</sup>IV | K132E (hom) |
| SIG-M5rNutlin<sup>20μM</sup>VI | R282W (het) |
| SIG-M5rNutlin<sup>20μM</sup>VIII | P27S (het) |
| SIG-M5rNutlin<sup>20μM</sup>IX | wild type |
| SIG-M5rNutlin<sup>20μM</sup>XI | c.196 del A (->Stop in Codon), V173L (het) |
| SIG-M5rNutlin<sup>20μM</sup>XV | wild type |
| SIG-M5rNutlin<sup>20μM</sup>XX | wild type |

<sup>1</sup> het, heterozygous; hom, homozygous
<sup>2</sup> All sub-lines share the identical mutation
<sup>3</sup> stop codon

**Figure 1.**
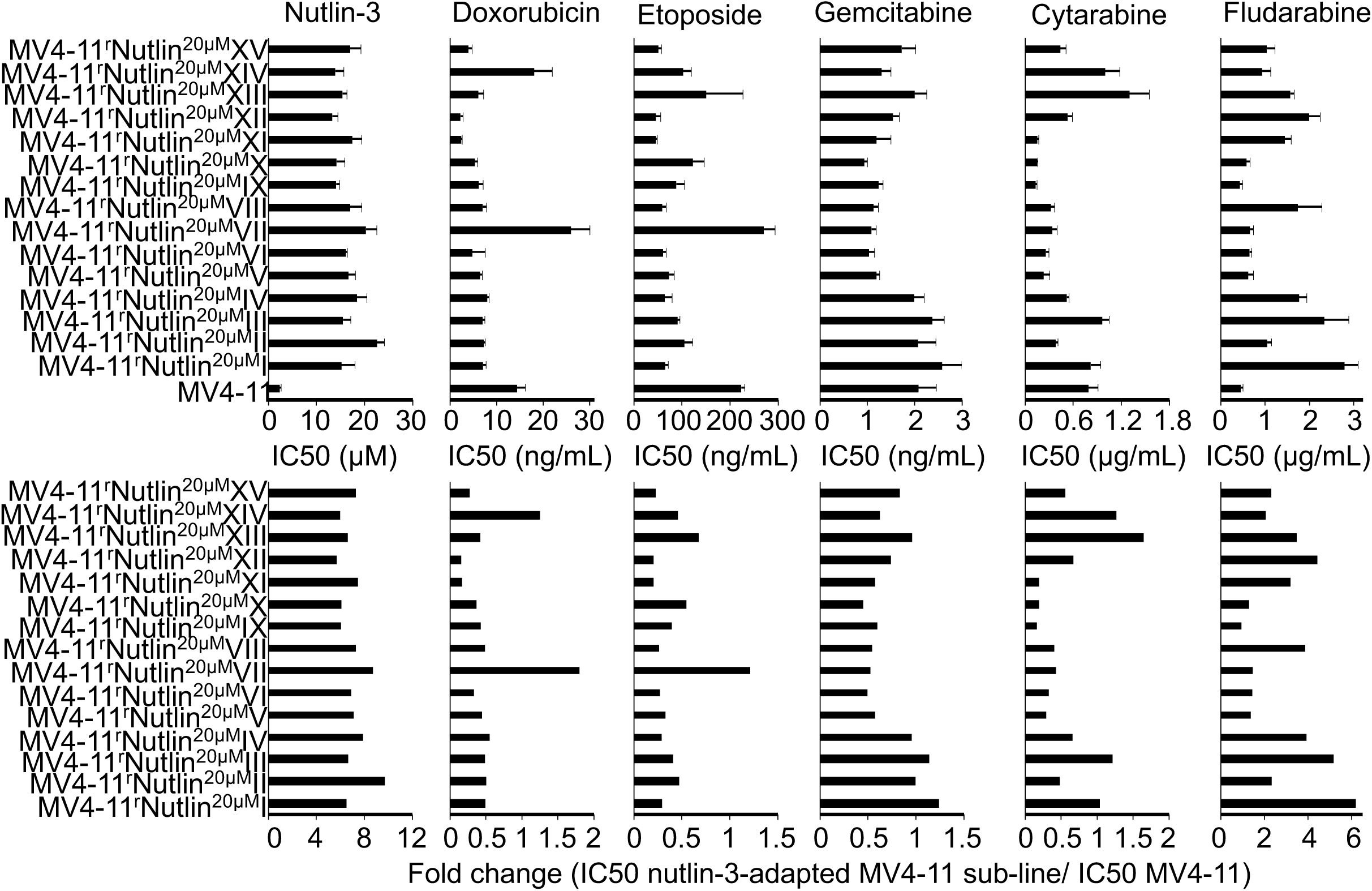
Drug sensitivity profiles of the AML cell line MV4-11 and its sub-lines adapted to nutlin-3 (20µM). Concentrations that inhibit cell viability by 50% (IC50) as determined by MTT assay after 120h incubation and relative sensitivity expressed as fold change (IC50 nutlin-3-resistant MV4-11 sub-line/ IC50 MV4-11). Numerical data are presented in Suppl. Table 1.

In the nutlin-3 adapted sub-lines of OCI-AML-2 (nutlin-3 IC50: 0.90µM), the nutlin-3 IC50s ranged from 14.8µM (OCI-AML-2^r^Nutlin^20µM^XI, resistance factor: 16.4) to 19.9µM (OCI-AML-2^r^Nutlin^20µM^II, resistance factor: 22.1) (Figure 2, Suppl. Table 1). In the OCI-AML-3 (nutlin-3 IC50: 1.75µM) sub-lines, the nutlin-3 IC50s ranged from 11.3µM (OCI-AML-3^r^Nutlin^20µM^XII, resistance factor: 6.5) to 20.62µM (OCI-AML-3^r^Nutlin^20µM^XI, resistance factor 11.8) (Figure 3, Suppl. Table 1) and in the SIG-M5 (Nutlin-3 IC50: 1.27µM) sub-lines from 3.64µM (SIG-M5^r^Nutlin^20µM^XV, resistance factor: 2.9) to 23.5µM (SIG-M5^r^Nutlin^20µM^XI, resistance factor: 18.5) (Figure 4, Suppl. Table 1).

**Figure 2.**
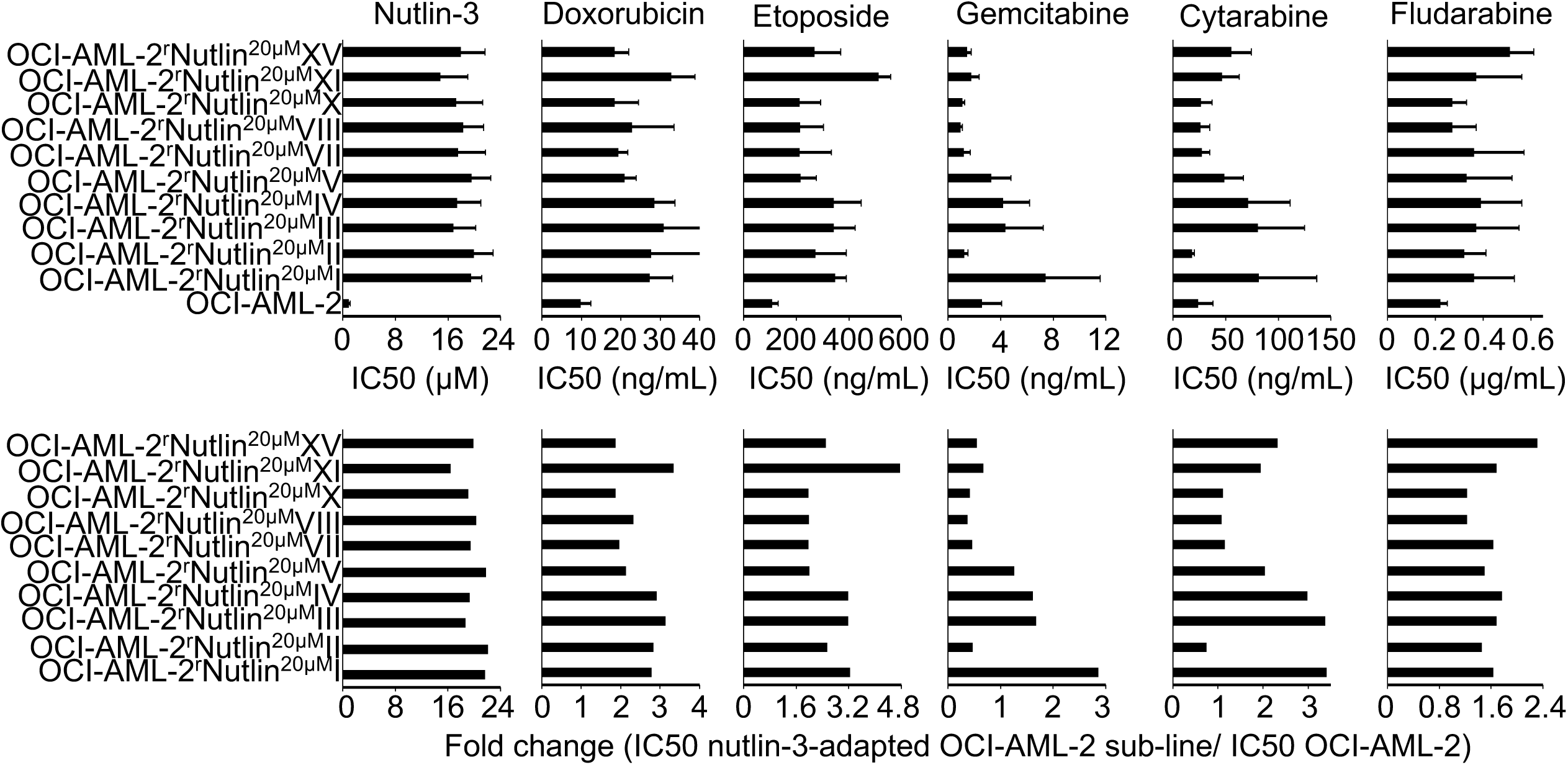
Drug sensitivity profiles of the AML cell line OCI-AML-2 and its sub-lines adapted to nutlin-3 (20µM). Concentrations that inhibit cell viability by 50% (IC50) as determined by MTT assay after 120h incubation and relative sensitivity expressed as fold change (IC50 nutlin-3-resistant OCI-AML-2 sub-line/ IC50 OCI-AML-2). Numerical data are presented in Suppl. Table 1.

**Figure 3.**
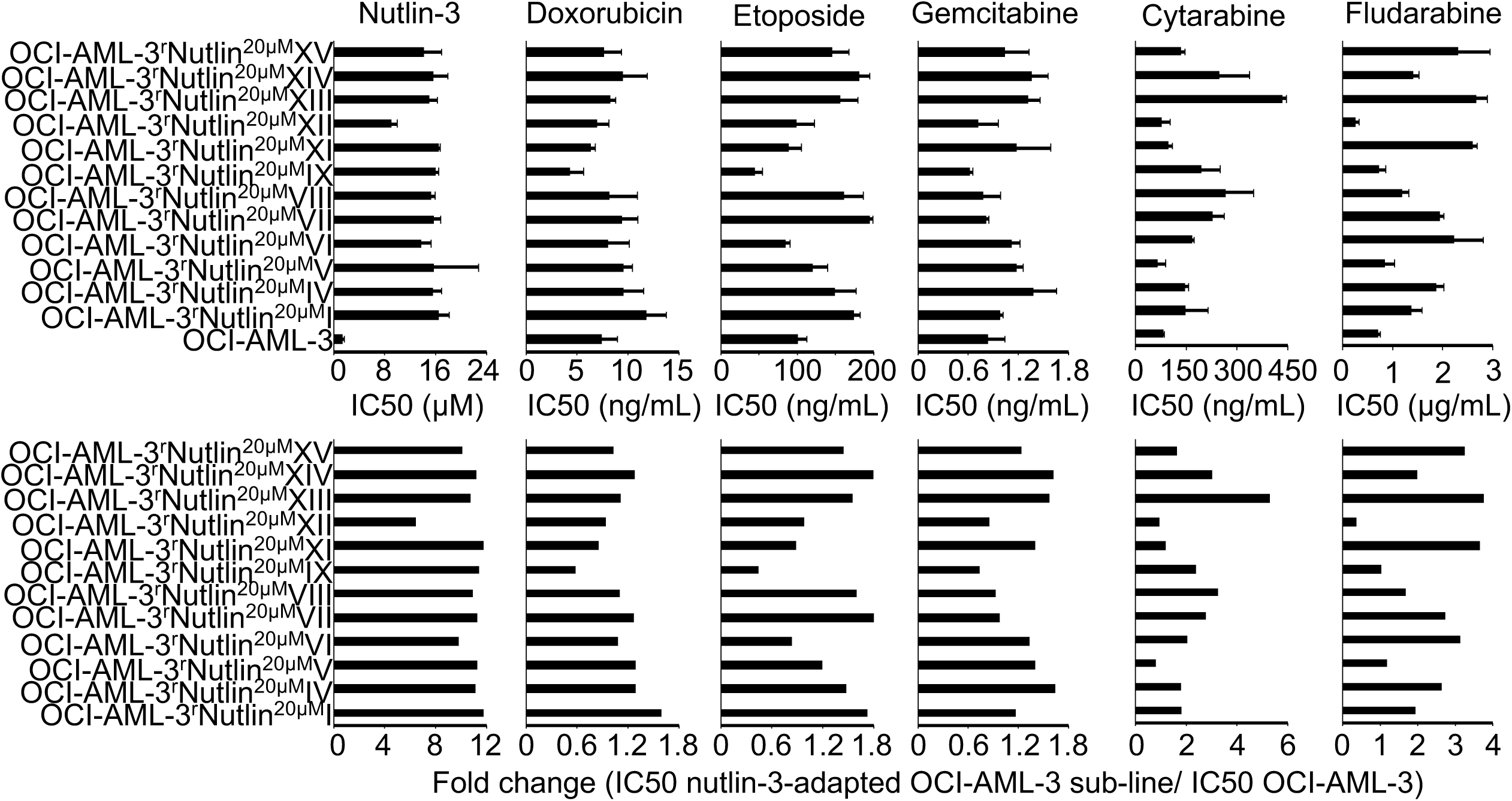
Drug sensitivity profiles of the AML cell line OCI-AML-3 and its sub-lines adapted to nutlin-3 (20µM). Concentrations that inhibit cell viability by 50% (IC50) as determined by MTT assay after 120h incubation and relative sensitivity expressed as fold change (IC50 nutlin-3-resistant OCI-AML-3 sub-line/ IC50 OCI-AML-3). Numerical data are presented in Suppl. Table 1.

**Figure 4.**
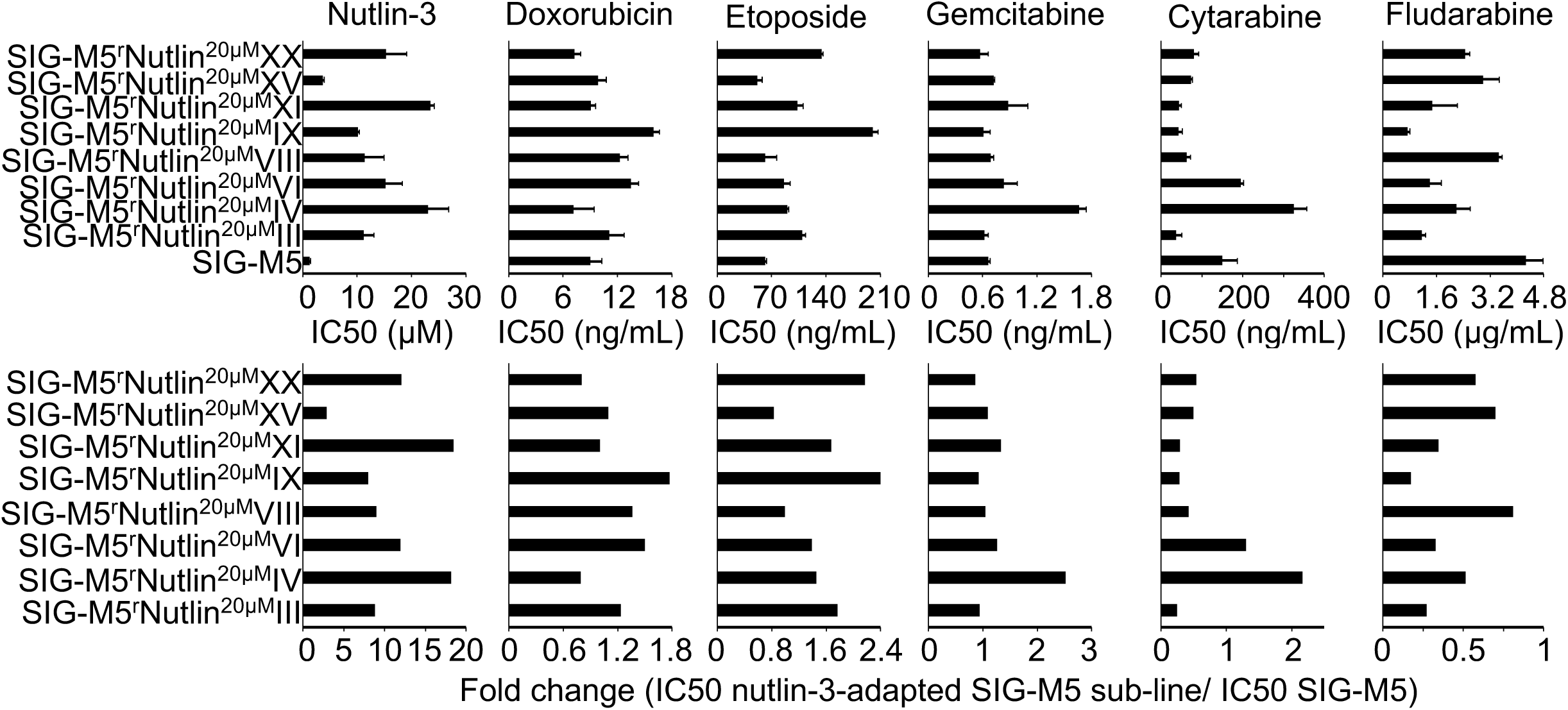
Drug sensitivity profiles of the AML cell line SIG-M5 and its sub-lines adapted to nutlin-3 (20µM). Concentrations that inhibit cell viability by 50% (IC50) as determined by MTT assay after 120h incubation and relative sensitivity expressed as fold change (IC50 nutlin-3-resistant SIG-M5 sub-line/ IC50 SIG-M5). Numerical data are presented in Suppl. Table 1.

### *TP53* status of nutlin-3-adapted AML cell lines and nutlin-3 resistance

The determination of the *TP53* status in the nutlin-3-adapted AML sub-lines revealed that all MV4-11 sub-lines harboured the same heterozygous R248W mutation and that all OCI-AML-2 sub-lines harboured the same heterozygous Y220C mutation (Table 1). In contrast, the OCI-AML-3 and SIG-M5 sub-lines harboured a range of different *TP53* mutations and included sub-lines that had retained wild-type *TP53*. (Table 1). In concordance, 219 out of 12418 reads of the appropriate *TP53* region in the parental MV4-11 cell line indicated the presence of alleles with an R248W mutation and 98 out of 907 reads indicated the presence of alleles with a Y220C mutation in the parental OCI-AML-2 cell line. In contrast, the mutations detected in the nutlin-3-adapted OCI-AML-3-and SIG-M5-sub-lines could not be detected in the respective parental cell lines. Also, MV4-11 and OCI-AML-2 could be adapted to nutlin-3 in 12-15 passages, whereas the nutlin-3 adaptation of OCI-AML-3 and SIG-M5 required 30-35 passages. This indicates that MV4-11 and OCI-AML-2 contain pre-existing *TP53*-mutant subpopulation that are selected by nutlin-3 treatment, while nutlin-3 treatment resulted in *de novo TP53* mutations in OCI-AML-3 and SIG-M5. These results are consistent with those obtained from other cancer entities [35,36,39,41].

Most of the *TP53* mutations are in the DNA binding domain (aa 102-292). The R248W mutation in the nutlin-3-adapted MV4-11 sub-lines and the Y220C mutation in the nutlin-3-adapted OCI-AML-2 sub-lines belong to the ten most commonly mutated *TP53* positions. 12 of the further 13 mutations are also located in the DNA binding domain and are known or expected to affect p53 function. Codon 27 is located in the transactivation domain, which is relevant for the MDM2-p53 interaction. The P27S mutation is known to increase the binding affinity of p53 to MDM2 [50-53].

There was no obvious relationship between the nutlin-3 IC50 in the parental cell lines in which nutlin-3 selected pre-existing *TP53*-mutant subpopulations (MV4-11: 2.33µM, OCI-AML-2: 0.90µM) and those parental cell lines in which nutlin-3 induced *de novo TP53*-mutations (OCI-AML-3: 1.75µM, SIG-M5: 1.27µM). The nutlin-3-adapted sub-lines displayed similar nutlin-3 IC50s independently of the mechanism of resistance formation or nutlin-3 sensitivity of the respective parental cell line (Figure 5). The fold changes (nutlin-3 IC50 resistant sub-line/ nutlin-3 IC50 respective parental cell line) were typically higher in parental cell lines that displayed lower nutlin-3 IC50 values (Figure 5). In the OCI-AML-3-and SIG-M5-sub-lines, there was no significant difference between the nutlin-3 IC50s in the *TP53*-mutant and *TP53* wild-type cell lines (Figure 5).

**Figure 5.**
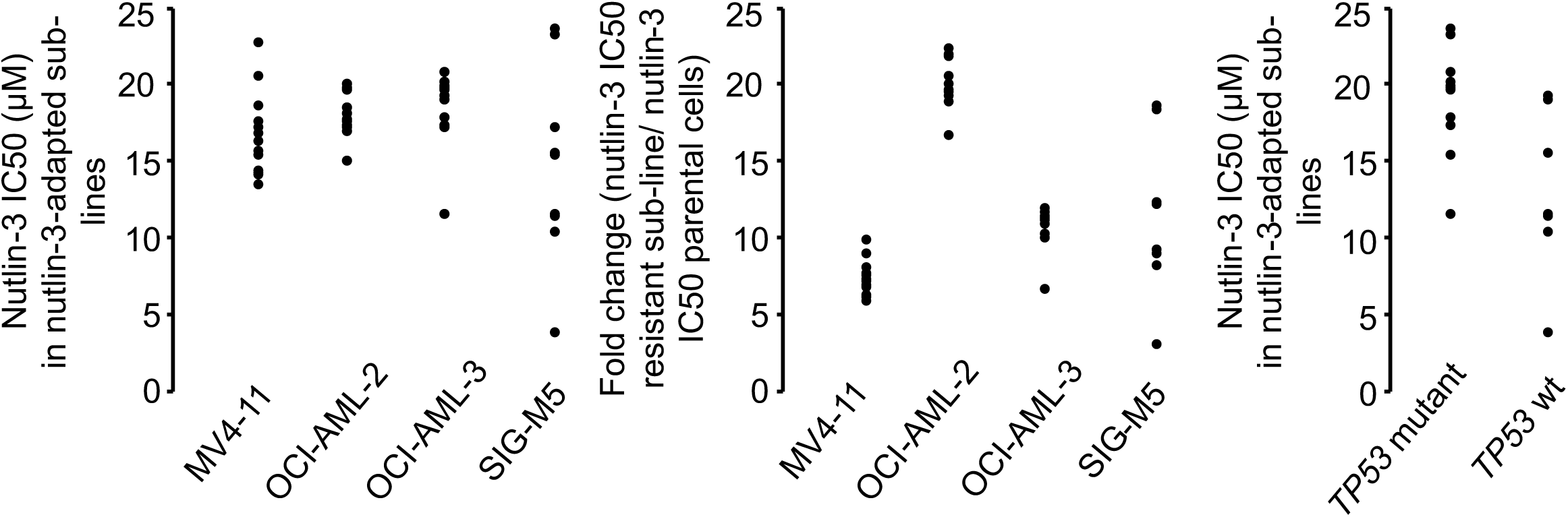
Distribution of the nutlin-3 IC50 values in the nutlin-3-adapted AML sub-lines. The IC50 values are presented as they are and as fold changes (nutlin-3 IC50 nutlin-3-adapted sub-line/ nutlin-3 IC50 respective parental cell line). In addition, the distribution of the nutlin-3 IC50 values is presented in the nutlin-3-adapted OCI-AML-3- and SIG-M5-sub-lines in dependence of their *TP53* mutation status. Numerical data are presented in Suppl. Table 1.

### Cross-resistance profiles in the nutlin-3-adapted AML sub-lines

Next, we determined sensitivity profiles of the nutlin-3-adapted AML sub-lines to doxorubicin, etoposide, gemcitabine, cytarabine, and fludarabine (Figure 1-4, Suppl. Table 1). According to the relative sensitivity of the nutlin-3-adapted sub-lines relative to the respective parental cell lines, sub-lines were categorised as more sensitive (IC50 nutlin-3-adapted sub-line/ IC50 respective parental cell line <0.5), less sensitive (IC50 nutlin-3-adapted sub-line/ IC50 respective parental cell line >2), or similarly sensitive (IC50 nutlin-3-adapted sub-line/ IC50 respective parental cell line >0.5 and <2) (Figure 6).

**Figure 6.**
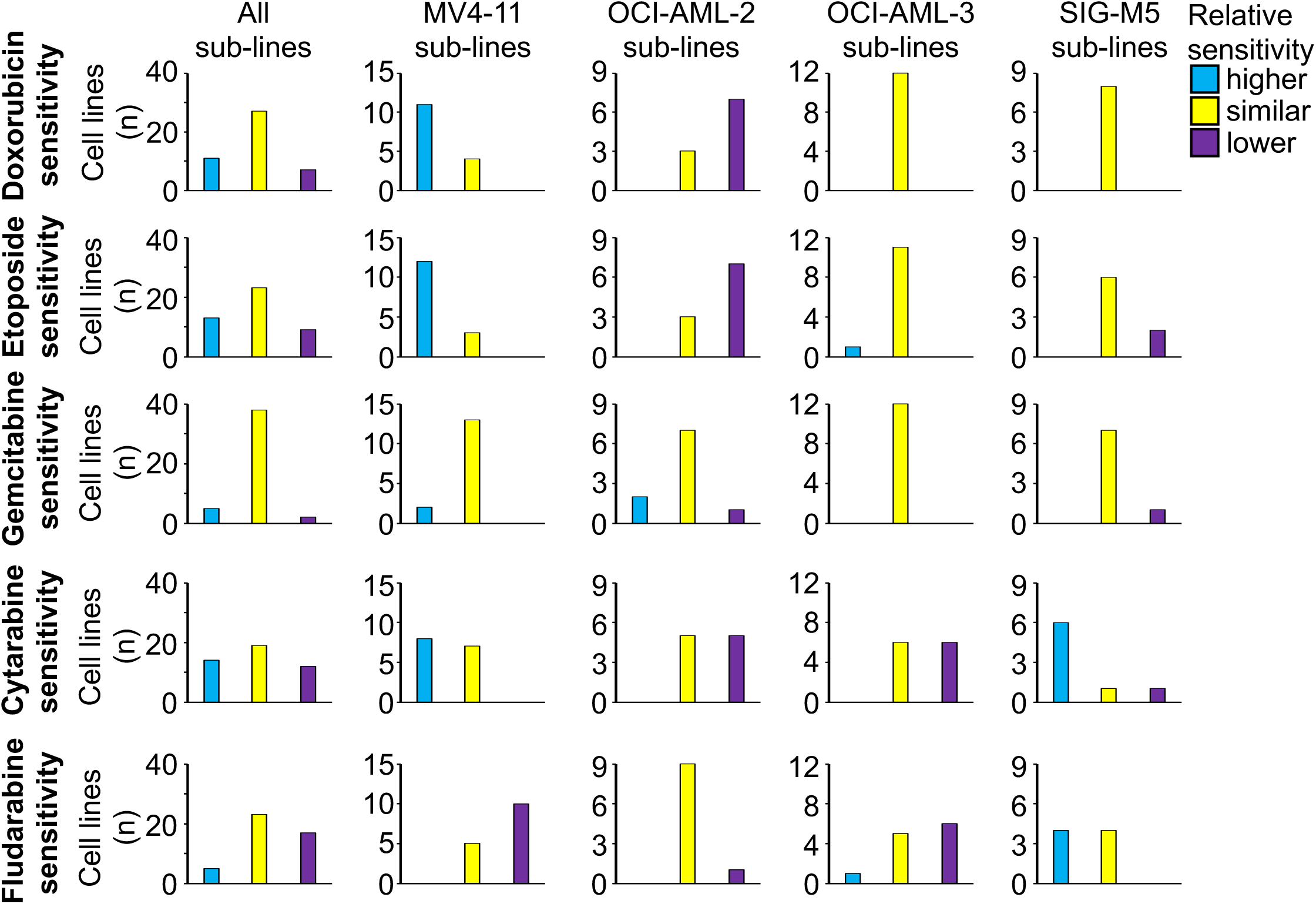
Nutlin-3-adapted AML sub-lines that display decreased, similar, or increased sensitivity to doxorubicin, etoposide, gemcitabine, cytarabine, or fludarabine relative to the respective parental cell lines. The nutlin-3-adapted AML sub-lines were categorised as cell lines that display a higher drug sensitivity than the respective parental cell line (IC50 nutlin-3-adapted sub-line/ IC50 respective parental cell line <0.5, blue bars), a similar drug sensitivity as the respective parental cell line (IC50 nutlin-3-adapted sub-line/ IC50 respective parental cell line >0.5 and <2, yellow bars), or a lower drug sensitivity than the respective parental cell line (IC50 nutlin-3-adapted sub-line/ IC50 respective parental cell line >2, purple bars). Numerical data are presented in Suppl. Table 1.

The sensitivity profiles indicated drug-and cell line-specific differences. Nutlin-3-resistance was not generally associated with increased resistance to other drugs (Figure 6). There was a noticeable heterogeneity in the drug response within the nutlin-3-resistant sub-lines of each parental cell line (Figure 1-4, 7). This included the MV4-11 and OCI-AML-2 sub-lines, although nutlin-3 had selected pre-existing *TP53-*mutant subpopulations in them. The maximum fold difference between nutlin-3-adapted sub-lines of the same parental cell line was 11.4 with MV4-11^r^Nutlin^20µM^XII having a doxorubicin IC50 of 2.28ng/mL and MV4-11^r^Nutlin^20µM^VII having a doxorubicin IC50 of 26.0ng/mL (Figure 7).

**Figure 7.**
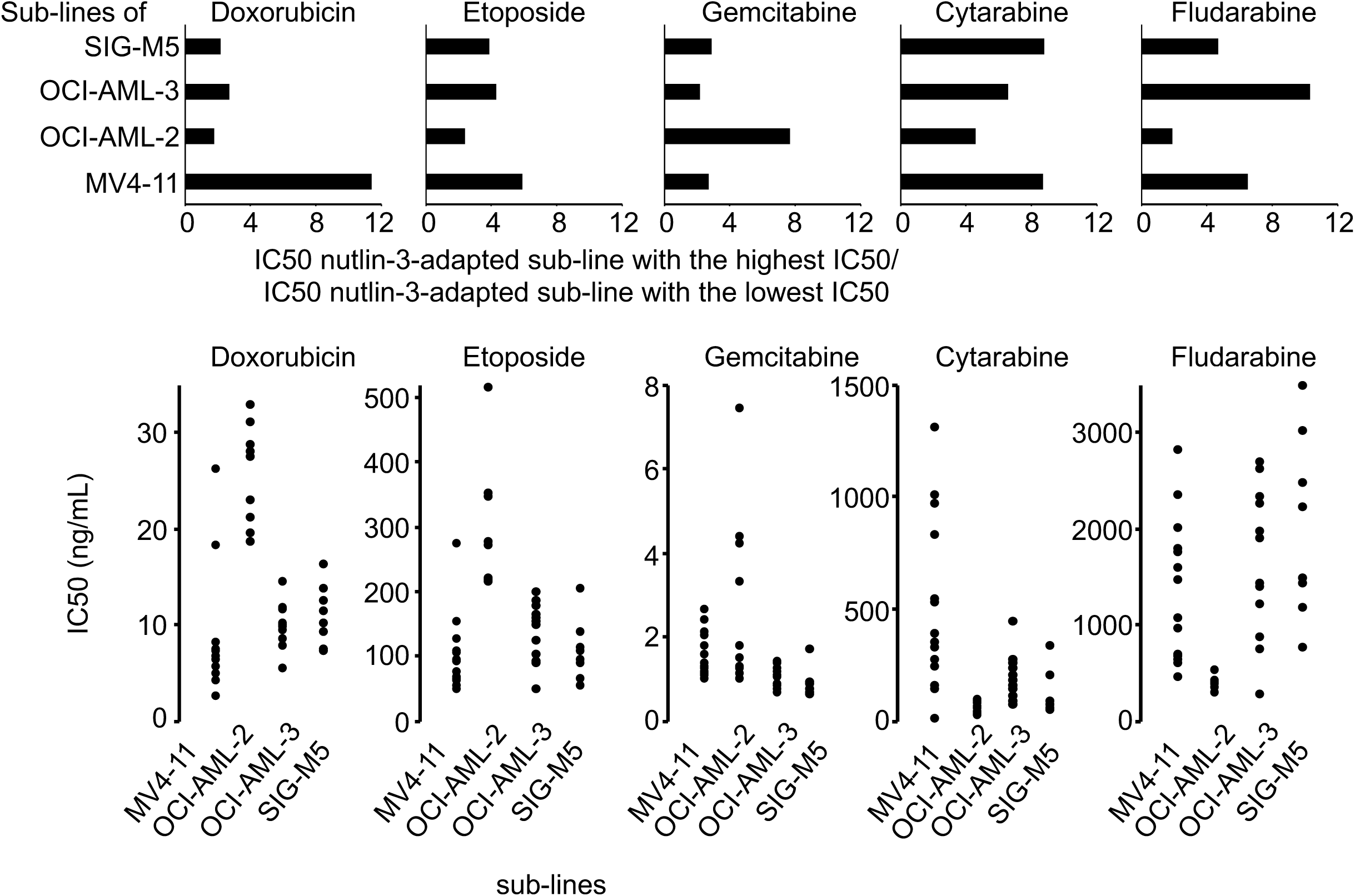
Comparison of the response of individual nutlin-3-adapted AML sub-lines to doxorubicin, etoposide, gemcitabine, cytarabine, or fludarabine. The fold change IC50 sub-line with the highest IC50/ IC50 sub-line with the lowest IC50 are presented for each drug in the nutlin-3-adapted sub-lines of MV4-11, OCI-AML-2, OCI-AML-3, and SIG-M5. In addition, the distribution of the IC50s of the individual cell lines are shown.

Finally, the drug response patterns were more similar between doxorubicin and etoposide than between these two drugs and the other agents (Figure 1-4, 6).

## Discussion

MDM2 inhibitors are currently being investigated in phase II and III clinical trials for AML (NCT02670044, NCT02545283). In various cell types, resistance formation to MDM2 inhibitors has previously been shown to be associated with the selection of pre-existing *TP53*-mutant cancer cell populations or the induction of *de novo TP53* mutations [3,35,36,39,41]. A clinical trial in liposarcoma patients confirmed that MDM2 inhibitor therapy is also associated with the emergence of *TP53* mutations in the clinic [40]. Here, we present a new set of models of acquired MDM2 inhibitor resistance in AML, in total 45 nutlin-3-adapted sub-lines of the AML cell lines MV4-11 (15 sub-lines), OCI-AML-2 (10 sub-lines), OCI-AML-3 (12 sub-lines), and SIG-M5 (8 sub-lines). Our results indicate that both mechanisms, selection of pre-existing *TP53*-mutant cancer cells and induction of *de novo TP53* mutations, are relevant in AML. Nutlin-3 consistently selected pre-existing *TP53*-mutant subpopulations in MV4-11 (R248W) and OCI-AML-2 (Y220C) cells. Interestingly, two other studies had also reported the emergence of R248W mutations in MV4-11 sub-lines. One study reported on an MDM2 inhibitor (SAR405838)-adapted MV4-11 sub-line with an R248W mutation [38]. Another one presented an R248W-mutant MV4-11 sub-line that had emerged during prolonged cell line cultivation [9]. This suggests the consistent presence of an MV4-11 subpopulation that harbours an R248W *TP53* mutation.

In contrast, the 12 nutlin-3-adapted OCI-AML-3 sub-lines included 9 *TP53*-mutant sub-lines, which all harboured different mutations, and 3 sub-lines that had retained wild-type *TP53*. Similarly, the 8 SIG-M5 sub-lines consisted of 4 *TP53*-mutant sub-lines, again each harbouring a different mutation, and 4 *TP53* wild-type sub-lines.

Notably, loss-of-p53-function has been associated with aggressive disease, chemoresistance, and dismal outcome in AML [54]. In patients with therapy-related AML, cytotoxic chemotherapy selected pre-existing *TP53*-mutant clones that were highly resistant to therapy [55,56]. However, resistance formation to nutlin-3 was not generally associated with cross-resistance to other anti-cancer drugs in AML cells. Hence, loss-of-p53-function does not always seem to mediate resistance to cytotoxic therapies directly. Indeed, RNAi-mediated depletion of p53 in SIG-M5 cells resulted in increased resistance to nutlin-3 but not to doxorubicin (Suppl. Figure 1). Notably, loss-of-p53 function may also indirectly increase the adaptability of AML cells to cytotoxic anti-cancer therapies, for example due to increased genomic instability [54].

In addition, the nutlin-3-adapted AML sub-lines displayed a noticeable heterogeneity in their responses to the anti-cancer drugs doxorubicin, etoposide, gemcitabine, cytarabine, and fludarabine. This also included the MV4-11 and OCI-AML-2 sub-lines, in which pre-existing *TP53*-mutant subpopulations had been selected by nutlin-3 treatment. Indeed, the highest fold change in the IC50 between the most sensitive and the most resistant nutlin-3-adapted sub-line of a given parental cell line was observed in MV4-11. The most doxorubicin-resistant MV4-11 sub-line (MV4-11^r^Nutlin^20µM^VII) displayed a doxorubicin IC50 of 26.0ng/mL, while the most doxorubicin-sensitive sub-line (MV4-11^r^Nutlin^20µM^XII) displayed a doxorubicin IC50 of 2.28ng/mL, resulting in an 11.4-fold difference. This indicates that the drug sensitivity profile of a nutlin-3-adapted AML subline cannot be predicted even if a defined pre-existing subpopulation of *TP53* mutant cells has been selected.

The doxorubicin and etoposide response profiles were more similar across the nutlin-3-adapted AML sub-lines than the sensitivity profiles of the other drugs. This may reflect a higher level of similarity between the mechanisms of action of doxorubicin and etoposide, which are both topoisomerase II inhibitors [57], compared to the other agents that are nucleoside analogues [58,59].

In conclusion, the investigation of 45 nutlin-3-adapted sub-lines of the AML cell lines MV4-11, OCI-AML-2, OCI-AML-3, and SIG-M5 showed that MDM2 inhibitors select, in dependence on the nature of a given AML cell population, pre-existing *TP53*-mutant subpopulations or induce *de novo TP53* mutations. Since MDM2 inhibitors are currently undergoing phase III clinical trials for the treatment of AML, patients should be monitored for the emergence of *TP53*-mutant leukaemia cells. The nutlin-3-adapted AML sub-lines showed a noticeable heterogeneity in their response to the cytotoxic anti-cancer drugs doxorubicin, etoposide, gemcitabine, cytarabine, and fludarabine. This indicates that even if a given cancer cell population is repeatedly adapted to the same drug in independent experiments, each adaptation follows an individual process resulting in a subpopulation with unique features. A substantial heterogeneity in the drug response was even observed in the MV4-11 and OCI-AML-2 sub-lines, in which nutlin-3 had selected pre-existing *TP53*-mutant subpopulations. Hence, future individualised treatment protocols will depend on the detailed monitoring of the evolutionary processes in cancer cell populations in response to therapy and an in-depth understanding of the therapeutic implications of the observed changes.

### Abbreviation list

AML: acute myeloid leukaemia
IC50: concentration that inhibits cell viability by 50%
MTT: 3-(4,5-dimethylthiazol-2-yl)-2,5-diphenyltetrazolium bromide

## Ethics approval and consent to participate

Not applicable

### Consent for publication

Not applicable

## Availability of data and materials

All data generated or analysed during this study are included in this published article and its supplementary information files.

## Competing interests

The authors declare that they have no competing interests.

## Funding

The work was supported by the Hilfe für krebskranke Kinder Frankfurt e.V., the Frankfurter Stiftung für krebskranke Kinder, the Deutsche José Carreras Leukämie-Stiftung, and the Kent Cancer Trust. The funding bodies had no role in the design of the study, the collection, analysis, and interpretation of data, and in writing the manuscript.

## Authors’ contributions

All authors analysed data and read and approved the final manuscript. MMi and JCjr directed the study and wrote the manuscript. CS, FR, TR, and JCjr were involved in the generation of the nutlin-3-resistant cell lines and sensitivity testing. MMe, AN, and TS were involved in the *TP53* sequencing and analysed the resulting data together with MM, DS, and JCjr.

## Acknowledgement

Not applicable

---

**Suppl. Figure 1.**
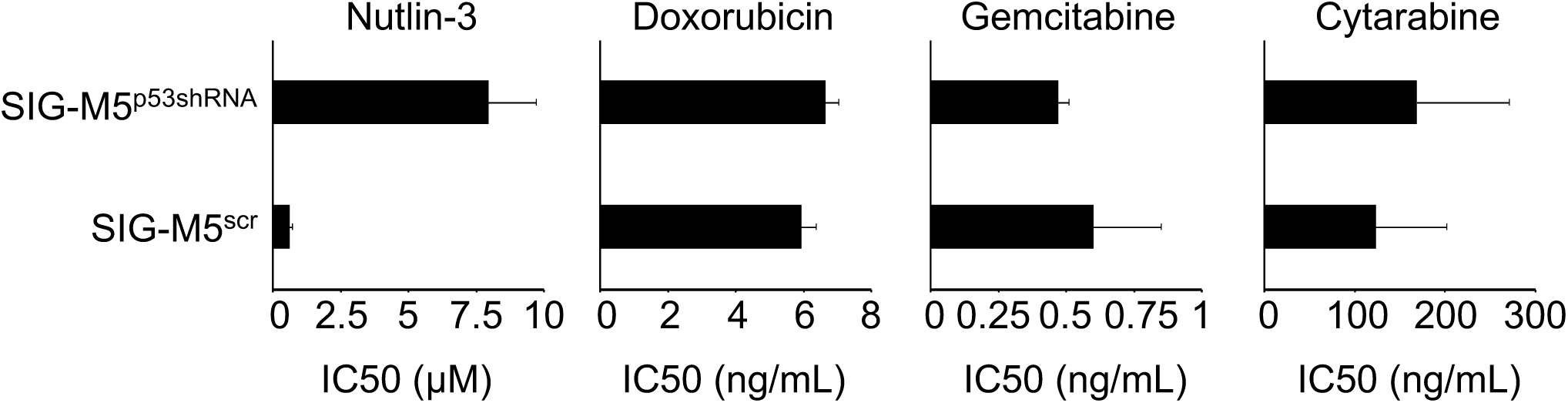
Drug sensitivity in SIG-M5 cells transduced with a lentiviral control vector encoding non-targeting (‘scrambled’) shRNA (SIG-M5^scr^) and SIG-M5 cells transduced with a lentiviral vector encoding shRNA targeting p53 (SIG-M5^p53shRNA^). Concentrations that reduce cell viability by 50% (IC50) were determined by MTT assay after 120h of incubation.

**Suppl. Table 1.** *TP53* status and drug sensitivity profiles in AML cell lines and their sub-lines adapted to nutlin-3 (20µM).

| Cell line | TP53 status | Drug concentration that reduces cell viability by 50% (IC50)<sup>1</sup> |  |  |  |  |  |
| --- | --- | --- | --- | --- | --- | --- | --- |
|  |  | nutlin-3<br>(μM) | doxorubicin<br>(ng/mL) | etoposide<br>(ng/mL) | gemcitabine<br>(ng/mL) | cytarabine<br>(μg/mL) | fludarabine<br>(μg/mL) |
| MV4-11 | wild type | 2.33 ± 0.35 | 14.4 ± 1.8 | 223 ± 7 | 2.08 ± 0.38 | 0.79 ± 0.12 | 0.45 ± 0.05 |
| MV4-11<sup>r</sup>Nutlin<sup>20μM</sup>I | R248W (het)<sup>2</sup> | 15.2 ± 2.8<br>(6.52)<sup>3</sup> | 7.15 ± 0.68<br>(0.49) | 65.3 ± 6.9<br>(0.29) | 2.58 ± 0.42<br>(1.24) | 0.82 ± 0.12<br>(1.04) | 2.79 ± 0.31<br>(6.20) |
| MV4-11<sup>r</sup>Nutlin<sup>20μM</sup>II | R248W (het) | 22.6 ± 1.5<br>(9.70) | 7.23 ± 0.36<br>(0.50) | 106 ± 17<br>(0.47) | 2.07 ± 0.38<br>(1.00) | 0.38 ± 0.03<br>(0.48) | 1.05 ± 0.10<br>(2.33) |
| MV4-11<sup>r</sup>Nutlin<sup>20μM</sup>III | R248W (het) | 15.5 ± 1.6<br>(6.65) | 6.99 ± 0.56<br>(0.48) | 91.2 ± 4.9<br>(0.41) | 2.37 ± 0.26<br>(1.14) | 0.96 ± 0.09<br>(1.22) | 2.33 ± 0.56<br>(5.18) |
| MV4-11<sup>r</sup>Nutlin<sup>20μM</sup>IV | R248W (het) | 18.4 ± 2.1<br>(7.90) | 8.00 ± 0.42<br>(0.55) | 64.3 ± 15.5<br>(0.29) | 1.99 ± 0.21<br>(0.96) | 0.52 ± 0.03<br>(0.66) | 1.77 ± 0.17<br>(3.93) |
| MV4-11<sup>r</sup>Nutlin<sup>20μM</sup>V | R248W (het) | 16.6 ± 1.5<br>(7.12) | 6.46 ± 0.52<br>(0.45) | 73.5 ± 10.9<br>(0.33) | 1.19 ± 0.07<br>(0.57) | 0.23 ± 0.08<br>(0.29) | 0.62 ± 0.12<br>(1.38) |
| MV4-11<sup>r</sup>Nutlin<sup>20μM</sup>VI | R248W (het) | 16.1 ± 0.3<br>(6.91) | 4.77 ± 2.85<br>(0.33) | 60.5 ± 7.3<br>(0.27) | 1.03 ± 0.12<br>(0.49) | 0.26 ± 0.04<br>(0.33) | 0.65 ± 0.05<br>(1.44) |
| MV4-11<sup>r</sup>Nutlin<sup>20μM</sup>VII | R248W (het) | 20.3 ± 2.2<br>(8.71) | 26.0 ± 4.1<br>(1.80) | 271 ± 23<br>(1.21) | 1.09 ± 0.10<br>(0.52) | 0.34 ± 0.06<br>(0.43) | 0.66 ± 0.08<br>(1.47) |
| MV4-11<sup>r</sup>Nutlin<sup>20μM</sup>VIII | R248W (het) | 17.0 ± 2.4<br>(7.30) | 7.05 ± 0.84<br>(0.49) | 59.3 ± 7.8<br>(0.27) | 1.13 ± 0.10<br>(0.54) | 0.32 ± 0.05<br>(0.41) | 1.74 ± 0.54<br>(3.87) |
| MV4-11<sup>r</sup>Nutlin<sup>20μM</sup>IX | R248W (het) | 14.1 ± 0.7<br>(6.05) | 6.18 ± 0.96<br>(0.43) | 88.7 ± 17.1<br>(0.40) | 1.24 ± 0.09<br>(0.60) | 0.13 ± 0.02<br>(0.16) | 0.43 ± 0.06<br>(0.96) |
| MV4-11<sup>r</sup>Nutlin<sup>20μM</sup>X | R248W (het) | 14.2 ± 1.7<br>(6.09) | 5.33 ± 0.65<br>(0.37) | 123 ± 24<br>(0.55) | 0.94 ± 0.06<br>(0.45) | 0.15 ± 0.01<br>(0.19) | 0.58 ± 0.08<br>(1.29) |
| MV4-11<sup>r</sup>Nutlin<sup>20μM</sup>XI | R248W (het) | 17.4 ± 2.0<br>(7.47) | 2.42 ± 0.27<br>(0.17) | 46.1 ± 3.1<br>(0.21) | 1.19 ± 0.31<br>(0.57) | 0.15 ± 0.02<br>(0.19) | 1.44 ± 0.15<br>(3.20) |
| MV4-11<sup>r</sup>Nutlin<sup>20μM</sup>XII | R248W (het) | 13.3 ± 1.2<br>(5.71) | 2.28 ± 0.56<br>(0.16) | 46.1 ± 9.7<br>(0.21) | 1.54 ± 0.14<br>(0.74) | 0.53 ± 0.06<br>(0.67) | 1.99 ± 0.25<br>(4.42) |
| MV4-11<sup>r</sup>Nutlin<sup>20μM</sup>XIII | R248W (het) | 15.4 ± 0.9<br>(6.61) | 6.13 ± 0.48<br>(0.42) | 151 ± 76<br>0.68 | 2.00 ± 0.25<br>(0.96) | 1.30 ± 0.25<br>(1.65) | 1.57 ± 0.09<br>(3.49) |
| MV4-11<sup>r</sup>Nutlin<sup>20μM</sup>XIV | R248W (het) | 13.9 ± 1.8<br>(5.97) | 18.1 ± 3.9<br>(1.25) | 102 ± 17<br>(0.46) | 1.30 ± 0.20<br>(0.63) | 1.00 ± 0.18<br>(1.27) | 0.93 ± 0.20<br>(2.07) |
| MV4-11<sup>r</sup>Nutlin<sup>20μM</sup>XV | R248W (het) | 17.0 ± 2.3<br>(7.30) | 3.95 ± 0.80<br>(0.27) | 51.3 ± 6.2<br>(0.23) | 1.73 ± 0.29<br>(0.83) | 0.44 ± 0.07<br>(0.56) | 1.04 ± 0.18<br>(2.31) |
|  |  | nutlin-3<br>(μM) | doxorubicin<br>(ng/mL) | etoposide<br>(ng/mL) | gemcitabine<br>(ng/mL) | cytarabine<br>(ng/mL) | fludarabine<br>(μg/mL) |
| OCI-AML-2 | wild type | 0.90 ± 0.22 | 9.80 ± 2.61 | 107 ± 24 | 2.59 ± 1.48 | 23.9 ± 13.9 | 0.22 ± 0.03 |
| OCI-AML-2<sup>r</sup>Nutlin<sup>20μM</sup>I | Y220C (het) | 19.5 ± 1.6<br>(21.7) | 27.3 ± 5.7<br>(2.79) | 348 ± 41<br>(3.24) | 7.40 ± 4.16<br>(2.86) | 81.5 ± 55.1<br>(3.41) | 0.36 ± 0.17<br>(1.64) |
| OCI-AML-2<sup>r</sup>Nutlin<sup>20μM</sup>II | Y220C (het) | 19.9 ± 2.1<br>(22.1) | 27.7 ± 12.6<br>(2.83) | 273 ± 117<br>(2.54) | 1.23 ± 0.29<br>(0.47) | 17.8 ± 2.4<br>(0.74) | 0.32 ± 0.09<br>(1.45) |
| OCI-AML-2<sup>r</sup>Nutlin<sup>20μM</sup>III | Y220C (het) | 16.8 ± 3.4<br>(18.7) | 30.8 ± 9.5<br>(3.14) | 342 ± 83<br>(3.18) | 4.34 ± 2.88<br>(1.68) | 80.6 ± 44.2<br>(3.37) | 0.37 ± 0.18<br>(1.68) |
| OCI-AML-2<sup>r</sup>Nutlin<sup>20μM</sup>IV | Y220C (het) | 17.4 ± 3.6<br>(19.3) | 28.5 ± 5.2<br>(2.91) | 342 ± 104<br>(3.18) | 4.18 ± 2.01<br>(1.61) | 71.2 ± 40.0<br>(2.98) | 0.39 ± 0.17<br>(1.77) |
| OCI-AML-2<sup>r</sup>Nutlin<sup>20μM</sup>V | Y220C (het) | 19.6 ± 2.9<br>(21.8) | 20.9 ± 2.9<br>(2.13) | 216 ± 61<br>(2.01) | 3.27 ± 1.51<br>(1.26) | 48.7 ± 18.0<br>(2.04) | 0.33 ± 0.19<br>(1.50) |
| OCI-AML-2<sup>r</sup>Nutlin<sup>20μM</sup>VII | Y220C (het) | 17.5 ± 4.2<br>(19.4) | 19.3 ± 2.4<br>(1.97) | 212 ± 121<br>(1.97) | 1.21 ± 0.49<br>(0.47) | 27.4 ± 7.5<br>(1.15) | 0.36 ± 0.21<br>(1.64) |
| OCI-AML-2<sup>r</sup>Nutlin<sup>20μM</sup>VIII | Y220C (het) | 18.3 ± 3.1<br>(20.3) | 22.8 ± 10.7<br>(2.33) | 214 ± 90<br>(1.99) | 0.96 ± 0.14<br>(0.37) | 25.8 ± 9.1<br>(1.08) | 0.27 ± 0.10<br>(1.23) |
| OCI-AML-2<sup>r</sup>Nutlin<sup>20μM</sup>X | Y220C (het) | 17.2 ± 4.1<br>(19.1) | 18.4 ± 6.1<br>(1.88) | 212 ± 80<br>(1.97) | 1.08 ± 0.20<br>(0.42) | 26.6 ± 10.5<br>(1.11) | 0.27 ± 0.06<br>(1.23) |
| OCI-AML-2<sup>r</sup>Nutlin<sup>20μM</sup>XI | Y220C (het) | 14.8 ± 4.2<br>(16.4) | 32.7 ± 6.1<br>(3.34) | 511 ± 47<br>(4.76) | 1.75 ± 0.63<br>(0.68) | 46.5 ± 16.2<br>(1.95) | 0.37 ± 0.19<br>(1.68) |
| OCI-AML-2<sup>r</sup>Nutlin<sup>20μM</sup>XV | Y220C (het) | 17.9 ± 3.7<br>(19.9) | 18.4 ± 3.66<br>(1.88) | 268 ± 101<br>(2.50) | 1.43 ± 0.33<br>(0.55) | 55.5 ± 18.7<br>(2.32) | 0.51 ± 0.10<br>(2.32) |
|  |  | nutlin-3<br>(μM) | doxorubicin<br>(ng/mL) | etoposide<br>(ng/mL) | gemcitabine<br>(ng/mL) | cytarabine<br>(ng/mL) | fludarabine<br>(μg/mL) |
| OCI-AML-3 | wild type | 1.75 ± 0.30 | 8.90 ± 1.89 | 101 ± 12 | 0.84 ± 0.20 | 82.1 ± 3.4 | 0.71 ± 0.04 |
| OCI-AML-3<sup>r</sup>Nutlin<sup>20μM</sup>I | R196*4 (hom) | 20.6 ± 2.1<br>(11.7) | 14.2 ± 2.4<br>(1.60) | 174 ± 9<br>(1.73) | 0.98 ± 0.04<br>(1.17) | 148 ± 66<br>(1.80) | 1.38 ± 0.21<br>(1.94) |
| OCI-AML-3<sup>r</sup>Nutlin<sup>20μM</sup>IV | R273S (het) | 19.5 ± 1.6<br>(11.1) | 11.5 ± 2.4<br>(1.29) | 149 ± 28<br>(1.48) | 1.38 ± 0.28<br>(1.64) | 147 ± 10<br>(1.49) | 1.87 ± 0.15<br>(2.63) |
| OCI-AML-3<sup>r</sup>Nutlin<sup>20μM</sup>V | S215G (het) | 19.7 ± 8.7<br>(11.3) | 11.5 ± 1.0<br>(1.29) | 121 ± 19<br>(1.20) | 1.18 ± 0.08<br>(1.40) | 65.3 ± 23.4<br>(0.80) | 0.84 ± 0.20<br>(1.18) |
| OCI-AML-3<sup>r</sup>Nutlin<sup>20μM</sup>VI | C176F (het) | 17.2 ± 1.9<br>(9.83) | 9.65 ± 2.51<br>(1.08) | 84.8 ± 5.8<br>(0.84) | 1.12 ± 0.10<br>(1.33) | 168 ± 5<br>(2.05) | 2.23 ± 0.58<br>(3.14) |
| OCI-AML-3<sup>r</sup>Nutlin<sup>20μM</sup>VII | G244S (het) | 19.7 ± 1.3<br>(11.3) | 11.3 ± 1.9<br>(1.27) | 195 ± 4<br>(1.94) | 0.82 ± 0.03<br>(0.98) | 228 ± 34<br>(2.78) | 1.94 ± 0.08<br>(2.73) |
| OCI-AML-3<sup>r</sup>Nutlin<sup>20μM</sup>VIII | wild-type | 19.1 ± 0.9<br>(10.9) | 9.83 ± 3.29<br>(1.10) | 161 ± 26<br>(1.60) | 0.78 ± 0.21<br>(0.93) | 266 ± 83<br>(3.24) | 1.19 ± 0.13<br>(1.68) |
| OCI-AML-3<sup>r</sup>Nutlin<sup>20μM</sup>IX | c.485 del 6bp<br>(TCTACA)<br>het,<br>IYK->K<br>(p.162..p.164) | 20.0 ± 0.6<br>(11.4) | 5.19 ± 1.61<br>(0.58) | 44.9 ± 9.6<br>(0.45) | 0.62 ± 0.04<br>(0.74) | 195 ± 55<br>(2.38) | 0.73 ± 0.13<br>(1.03) |
| OCI-AML-3<sup>r</sup>Nutlin<sup>20μM</sup>XI | G266V (het) | 20.6 ± 0.3<br>(11.8) | 7.63 ± 0.55<br>(0.86) | 89.3 ± 16.3<br>(0.89) | 1.18 ± 0.40<br>(1.40) | 97.0 ± 11.8<br>(1.18) | 2.60 ± 0.09<br>(3.66) |
| OCI-AML-3<sup>r</sup>Nutlin<sup>20μM</sup>XII | wild type | 11.3 ± 1.2<br>(6.46) | 8.37 ± 1.42<br>(0.94) | 99 ± 24<br>(0.98) | 0.72 ± 0.24<br>(0.86) | 77.8 ± 25.2<br>(0.95) | 0.26 ± 0.07<br>(0.37) |
| OCI-AML-3<sup>r</sup>Nutlin<sup>20μM</sup>XIII | wild type | 18.8 ± 1.6<br>(10.7) | 9.91 ± 0.67<br>(1.11) | 157 ± 23<br>(1.55) | 1.32 ± 0.14<br>(1.57) | 434 ± 12<br>(5.29) | 2.67 ± 0.22<br>(3.76) |
| OCI-AML-3<sup>r</sup>Nutlin<sup>20μM</sup>XIV | S215G (het) | 19.6 ± 2.8<br>(11.2) | 11.4 ± 2.9<br>(1.28) | 181 ± 14<br>(1.80) | 1.36 ± 0.20<br>(1.62) | 248 ± 89<br>(3.02) | 1.41 ± 0.12<br>(1.99) |
| OCI-AML-3<sup>r</sup>Nutlin<sup>20μM</sup>XV | R248Q (het) | 17.7 ± 3.5<br>(10.1) | 9.18 ± 2.10<br>(1.03) | 146 ± 22<br>(1.45) | 1.04 ± 0.29<br>(1.24) | 134 ± 13<br>(1.63) | 2.31 ± 0.64<br>(3.25) |

| Cell line | TP53 status | Drug concentration that reduces cell viability by 50% (IC<sub>50</sub>)<sup>1</sup> |  |  |  |  |  |
| --- | --- | --- | --- | --- | --- | --- | --- |
|  |  | nutlin-3<br>(μM) | doxorubicin<br>(ng/mL) | etoposide<br>(ng/mL) | gemcitabine<br>(ng/mL) | cytarabine<br>(ng/mL) | fludarabine<br>(μg/mL) |
| SIG-M5 | wild type | 1.27 ± 0.16 | 9.01 ± 1.26 | 61.9 ± 1.6 | 0.66 ± 0.02 | 150 ± 37 | 4.27 ± 0.52 |
| SIG-M5<sup>r</sup>Nutlin<sup>20μM</sup>III | wild type | 11.2 ± 1.9<br>(8.80) | 11 ± 1.6<br>(1.23) | 109 ± 4<br>(1.77) | 0.62 ± 0.04<br>(0.94) | 36.9 ± 13.7<br>(0.25) | 1.16 ± 0.11<br>(0.27) |
| SIG-M5<sup>r</sup>Nutlin<sup>20μM</sup>IV | K132E (hom) | 23.0 ± 3.8<br>(18.1) | 7.12 ± 2.31<br>(0.79) | 90.0 ± 2.3<br>(1.45) | 1.66 ± 0.08<br>(2.52) | 325 ± 32<br>(2.17) | 2.20 ± 0.41<br>(0.52) |
| SIG-M5<sup>r</sup>Nutlin<sup>20μM</sup>VI | R282W (het) | 15.2 ± 3.2<br>(11.9) | 13.5 ± 0.85<br>(1.50) | 86.2 ± 7.6<br>(1.39) | 0.83 ± 0.15<br>(1.26) | 195 ± 7<br>(1.30) | 1.40 ± 0.34<br>(0.33) |
| SIG-M5<sup>r</sup>Nutlin<sup>20μM</sup>VIII | P27S (het) | 11.4 ± 3.5<br>(8.98) | 12.3 ± 0.9<br>(1.36) | 61.6 ± 15.1<br>(1.00) | 0.69 ± 0.03<br>(1.05) | 63.0 ± 8.5<br>(0.42) | 3.46 ± 0.10<br>(0.81) |
| SIG-M5<sup>r</sup>Nutlin<sup>20μM</sup>IX | wild type | 10.1 ± 0.3<br>(7.97) | 16.0 ± 0.7<br>(1.77) | 200 ± 7<br>(3.23) | 0.61 ± 0.07<br>(0.92) | 42.5 ± 9.3<br>(0.28) | 0.74 ± 0.07<br>(0.17) |
| SIG-M5<sup>r</sup>Nutlin<sup>20μM</sup>XI | c.196 del A (->Stop in Codon), V173L (het) | 23.5 ± 0.7<br>(18.5) | 9.07 ± 0.54<br>(1.01) | 103 ± 7<br>(1.67) | 0.88 ± 0.22<br>(1.33) | 43.1 ± 6.6<br>(0.29) | 1.47 ± 0.75<br>(0.34) |
| SIG-M5<sup>r</sup>Nutlin<sup>20μM</sup>XV | wild type | 3.64 ± 0.29<br>(2.87) | 9.87 ± 0.90<br>(1.10) | 51.5 ± 6.4<br>(0.83) | 0.72 ± 0.01<br>(1.09) | 73.7 ± 3.4<br>(0.49) | 2.99 ± 0.48<br>(0.70) |
| SIG-M5<sup>r</sup>Nutlin<sup>20μM</sup>XX | wild type | 15.3 ± 3.8<br>(12.1) | 7.23 ± 0.71<br>(0.80) | 134 ± 2<br>(2.17) | 0.57 ± 0.09<br>(0.86) | 80.7 ± 11.1<br>(0.54) | 2.46 ± 0.13<br>(0.58) |

<sup>1</sup> Determined by MTT after a 120h incubation period<sup>2</sup> het, heterozygous; hom, homozygous
<sup>3</sup> Fold change (IC50 nutlin-3-adapted sub-line/ IC50 respective parental cell line) <sup>4</sup> Stop codon

## References

1. Tisato V, Voltan R, Gonelli A, Secchiero P, Zauli G. MDM2/X inhibitors under clinical evaluation: perspectives for the management of hematological malignancies and pediatric cancer. J Hematol Oncol. 2017;10:133.

2. Wade M, Li YC, Wahl GM. MDM2, MDMX and p53 in oncogenesis and cancer therapy. Nat Rev Cancer. 2013;13:83–96.

3. Cinatl J Jr, Speidel D, Hardcastle I, Michaelis M. Resistance acquisition to MDM2 inhibitors. Biochem Soc Trans. 2014;42:752–7.

4. Kojima K, Konopleva M, Samudio IJ, Shikami M, Cabreira-Hansen M, McQueen T, Ruvolo V, Tsao T, Zeng Z, Vassilev LT, Andreeff M. MDM2 antagonists induce p53-dependent apoptosis in AML: implications for leukemia therapy. Blood. 2005;106:3150–9.

5. Kojima K, Konopleva M, Samudio IJ, Schober WD, Bornmann WG, Andreeff M. Concomitant inhibition of MDM2 and Bcl-2 protein function synergistically induce mitochondrial apoptosis in AML. Cell Cycle. 2006;5:2778–86.

6. Secchiero P, Zerbinati C, Melloni E, Milani D, Campioni D, Fadda R, Tiribelli M, Zauli G. The MDM-2 antagonist nutlin-3 promotes the maturation of acute myeloid leukemic blasts. Neoplasia. 2007;9:853–61.

7. Kojima K, Konopleva M, Tsao T, Nakakuma H, Andreeff M. Concomitant inhibition of Mdm2-p53 interaction and Aurora kinases activates the p53-dependent postmitotic checkpoints and synergistically induces p53-mediated mitochondrial apoptosis along with reduced endoreduplication in acute myelogenous leukemia. Blood. 2008;112:2886–95.

8. Carter BZ, Mak DH, Schober WD, Koller E, Pinilla C, Vassilev LT, Reed JC, Andreeff M. Simultaneous activation of p53 and inhibition of XIAP enhance the activation of apoptosis signaling pathways in AML. Blood. 2010;115:306–14.

9. Kojima K, Konopleva M, Tsao T, Andreeff M, Ishida H, Shiotsu Y, Jin L, Tabe Y, Nakakuma H. Selective FLT3 inhibitor FI-700 neutralizes Mcl-1 and enhances p53-mediated apoptosis in AML cells with activating mutations of FLT3 through Mcl-1/Noxa axis. Leukemia. 2010;24:33–43.

10. Long J, Parkin B, Ouillette P, Bixby D, Shedden K, Erba H, Wang S, Malek SN. Multiple distinct molecular mechanisms influence sensitivity and resistance to MDM2 inhibitors in adult acute myelogenous leukemia. Blood. 2010;116:71–80.

11. Samudio IJ, Duvvuri S, Clise-Dwyer K, Watt JC, Mak D, Kantarjian H, Yang D, Ruvolo V, Borthakur G. Activation of p53 signaling by MI-63 induces apoptosis in acute myeloid leukemia cells. Leuk Lymphoma. 2010;51:911–9.

12. Thompson T, Andreeff M, Studzinski GP, Vassilev LT. 1,25-dihydroxyvitamin D3 enhances the apoptotic activity of MDM2 antagonist nutlin-3a in acute myeloid leukemia cells expressing wild-type p53. Mol Cancer Ther. 2010;9:1158–68.

13. Zhang W, Konopleva M, Burks JK, Dywer KC, Schober WD, Yang JY, McQueen TJ, Hung MC, Andreeff M. Blockade of mitogen-activated protein kinase/extracellular signal-regulated kinase kinase and murine double minute synergistically induces Apoptosis in acute myeloid leukemia via BH3-only proteins Puma and Bim. Cancer Res. 2010;70:2424–34.

14. McCormack E, Haaland I, Venås G, Forthun RB, Huseby S, Gausdal G, Knappskog S, Micklem DR, Lorens JB, Bruserud O, Gjertsen BT. Synergistic induction of p53 mediated apoptosis by valproic acid and nutlin-3 in acute myeloid leukemia. Leukemia. 2012;26:910–7.

15. Haaland I, Opsahl JA, Berven FS, Reikvam H, Fredly HK, Haugse R, Thiede B, McCormack E, Lain S, Bruserud O, Gjertsen BT. Molecular mechanisms of nutlin-3 involve acetylation of p53, histones and heat shock proteins in acute myeloid leukemia. Mol Cancer. 2014;13:116.

16. Weisberg E, Halilovic E, Cooke VG, Nonami A, Ren T, Sanda T, Simkin I, Yuan J, Antonakos B, Barys L, Ito M, Stone R, Galinsky I, Cowens K, Nelson E, Sattler M, Jeay S, Wuerthner JU, McDonough SM, Wiesmann M, Griffin JD. Inhibition of Wild-Type p53-Expressing AML by the Novel Small Molecule HDM2 Inhibitor CGM097. Mol Cancer Ther. 2015;14:2249–59.

17. Lehmann C, Friess T, Birzele F, Kiialainen A, Dangl M. Superior anti-tumor activity of the MDM2 antagonist idasanutlin and the Bcl-2 inhibitor venetoclax in p53 wild-type acute myeloid leukemia models. J Hematol Oncol. 2016;9:50.

18. Cassier PA, Castets M, Belhabri A, Vey N. Targeting apoptosis in acute myeloid leukaemia. Br J Cancer. 2017;117:1089–98.

19. Pan R, Ruvolo V, Mu H, Leverson JD, Nichols G, Reed JC, Konopleva M, Andreeff M. Synthetic Lethality of Combined Bcl-2 Inhibition and p53 Activation in AML: Mechanisms and Superior Antileukemic Efficacy. Cancer Cell. 2017;32:748-760.e6.

20. Seipel K, Marques MAT, Sidler C, Mueller BU, Pabst T. The Cellular p53 Inhibitor MDM2 and the Growth Factor Receptor FLT3 as Biomarkers for Treatment Responses to the MDM2-Inhibitor Idasanutlin and the MEK1 Inhibitor Cobimetinib in Acute Myeloid Leukemia. Cancers (Basel). 2018;10. pii: E170.

21. Andreeff M, Kelly KR, Yee K, Assouline S, Strair R, Popplewell L, Bowen D, Martinelli G, Drummond MW, Vyas P, Kirschbaum M, Iyer SP, Ruvolo V, González GM, Huang X, Chen G, Graves B, Blotner S, Bridge P, Jukofsky L, Middleton S, Reckner M, Rueger R, Zhi J, Nichols G, Kojima K. Results of the Phase I Trial of RG7112, a Small-Molecule MDM2 Antagonist in Leukemia. Clin Cancer Res. 2016;22:868–76.

22. Ravandi F, Gojo I, Patnaik MM, Minden MD, Kantarjian H, Johnson-Levonas AO, Fancourt C, Lam R, Jones MB, Knox CD, Rose S, Patel PS, Tibes R. A phase I trial of the human double minute 2 inhibitor (MK-8242) in patients with refractory/recurrent acute myelogenous leukemia (AML). Leuk Res. 2016 Sep;48:92-100.

23. Reis B, Jukofsky L, Chen G, Martinelli G, Zhong H, So WV, Dickinson MJ, Drummond M, Assouline S, Hashemyan M, Theron M, Blotner S, Lee JH, Kasner M, Yoon SS, Rueger R, Seiter K, Middleton SA, Kelly KR, Vey N, Yee K, Nichols G, Chen LC, Pierceall WE. Acute myeloid leukemia patients’ clinical response to idasanutlin (RG7388) is associated with pre-treatment MDM2 protein expression in leukemic blasts. Haematologica. 2016;101:e185–8.

24. Engelman JA, Zejnullahu K, Mitsudomi T, Song Y, Hyland C, Park JO, Lindeman N, Gale CM, Zhao X, Christensen J, Kosaka T, Holmes AJ, Rogers AM, Cappuzzo F, Mok T, Lee C, Johnson BE, Cantley LC, Jänne PA. MET amplification leads to gefitinib resistance in lung cancer by activating ERBB3 signaling. Science. 2007;316:1039–43.

25. Nazarian R, Shi H, Wang Q, Kong X, Koya RC, Lee H, Chen Z, Lee MK, Attar N, Sazegar H, Chodon T, Nelson SF, McArthur G, Sosman JA, Ribas A, Lo RS. Melanomas acquire resistance to B-RAF(V600E) inhibition by RTK or N-RAS upregulation. Nature. 2010;468:973–7.

26. Poulikakos PI, Persaud Y, Janakiraman M, Kong X, Ng C, Moriceau G, Shi H, Atefi M, Titz B, Gabay MT, Salton M, Dahlman KB, Tadi M, Wargo JA, Flaherty KT, Kelley MC, Misteli T, Chapman PB, Sosman JA, Graeber TG, Ribas A, Lo RS, Rosen N, Solit DB. RAF inhibitor resistance is mediated by dimerization of aberrantly spliced BRAF(V600E). Nature. 2011;480:387–90.

27. Domingo-Domenech J, Vidal SJ, Rodriguez-Bravo V, Castillo-Martin M, Quinn SA, Rodriguez-Barrueco R, Bonal DM, Charytonowicz E, Gladoun N, de la Iglesia-Vicente J, Petrylak DP, Benson MC, Silva JM, Cordon-Cardo C. Suppression of acquired docetaxel resistance in prostate cancer through depletion of notch-and hedgehog-dependent tumor-initiating cells. Cancer Cell. 2012;22:373–88.

28. Joseph JD, Lu N, Qian J, Sensintaffar J, Shao G, Brigham D, Moon M, Maneval EC, Chen I, Darimont B, Hager JH. A clinically relevant androgen receptor mutation confers resistance to second-generation antiandrogens enzalutamide and ARN-509. Cancer Discov. 2013;3:1020–9.

29. Korpal M, Korn JM, Gao X, Rakiec DP, Ruddy DA, Doshi S, Yuan J, Kovats SG, Kim S, Cooke VG, Monahan JE, Stegmeier F, Roberts TM, Sellers WR, Zhou W, Zhu P. An F876L mutation in androgen receptor confers genetic and phenotypic resistance to MDV3100 (enzalutamide). Cancer Discov. 2013;3:1030–43.

30. Crystal AS, Shaw AT, Sequist LV, Friboulet L, Niederst MJ, Lockerman EL, Frias RL, Gainor JF, Amzallag A, Greninger P, Lee D, Kalsy A, Gomez-Caraballo M, Elamine L, Howe E, Hur W, Lifshits E, Robinson HE, Katayama R, Faber AC, Awad MM, Ramaswamy S, Mino-Kenudson M, Iafrate AJ, Benes CH, Engelman JA. Patient-derived models of acquired resistance can identify effective drug combinations for cancer. Science. 2014;346:1480–6.

31. Niederst MJ, Sequist LV, Poirier JT, Mermel CH, Lockerman EL, Garcia AR, Katayama R, Costa C, Ross KN, Moran T, Howe E, Fulton LE, Mulvey HE, Bernardo LA, Mohamoud F, Miyoshi N, VanderLaan PA, Costa DB, Jänne PA, Borger DR, Ramaswamy S, Shioda T, Iafrate AJ, Getz G, Rudin CM, Mino-Kenudson M, Engelman JA. RB loss in resistant EGFR mutant lung adenocarcinomas that transform to small-cell lung cancer. Nat Commun. 2015;6:6377.

32. Göllner S, Oellerich T, Agrawal-Singh S, Schenk T, Klein HU, Rohde C, Pabst C, Sauer T, Lerdrup M, Tavor S, Stölzel F, Herold S, Ehninger G, Köhler G, Pan KT, Urlaub H, Serve H, Dugas M, Spiekermann K, Vick B, Jeremias I, Berdel WE, Hansen K, Zelent A, Wickenhauser C, Müller LP, Thiede C, Müller-Tidow C. Loss of the histone methyltransferase EZH2 induces resistance to multiple drugs in acute myeloid leukemia. Nat Med. 2017;23:69–78.

33. Schneider C, Oellerich T, Baldauf HM, Schwarz SM, Thomas D, Flick R, Bohnenberger H, Kaderali L, Stegmann L, Cremer A, Martin M, Lohmeyer J, Michaelis M, Hornung V, Schliemann C, Berdel WE, Hartmann W, Wardelmann E, Comoglio F, Hansmann ML, Yakunin AF, Geisslinger G, Ströbel P, Ferreirós N, Serve H, Keppler OT, Cinatl J Jr. SAMHD1 is a biomarker for cytarabine response and a therapeutic target in acute myeloid leukemia. Nat Med. 2017;23:250–5.

34. Aziz MH, Shen H, Maki CG. Acquisition of p53 mutations in response to the non-genotoxic p53 activator Nutlin-3. Oncogene. 2011 Nov 17;30(46):4678–86.

35. Michaelis M, Rothweiler F, Barth S, Cinatl J, van Rikxoort M, Löschmann N, Voges Y, Breitling R, von Deimling A, Rödel F, Weber K, Fehse B, Mack E, Stiewe T, Doerr HW, Speidel D, Cinatl J Jr. Adaptation of cancer cells from different entities to the MDM2 inhibitor nutlin-3 results in the emergence of p53-mutated multi-drug-resistant cancer cells. Cell Death Dis. 2011;2:e243.

36. Michaelis M, Rothweiler F, Agha B, Barth S, Voges Y, Löschmann N, von Deimling A, Breitling R, Doerr HW, Rödel F, Speidel D, Cinatl J Jr. Human neuroblastoma cells with acquired resistance to the p53 activator RITA retain functional p53 and sensitivity to other p53 activating agents. Cell Death Dis. 2012;3:e294.

37. Jones RJ, Bjorklund CC, Baladandayuthapani V, Kuhn DJ, Orlowski RZ. Drug resistance to inhibitors of the human double minute-2 E3 ligase is mediated by point mutations of p53, but can be overcome with the p53 targeting agent RITA. Mol Cancer Ther. 2012;11:2243–53.

38. Hoffman-Luca CG, Ziazadeh D, McEachern D, Zhao Y, Sun W, Debussche L, Wang S. Elucidation of Acquired Resistance to Bcl-2 and MDM2 Inhibitors in Acute Leukemia In Vitro and In Vivo. Clin Cancer Res. 2015;21:2558–68.

39. Drummond CJ, Esfandiari A, Liu J, Lu X, Hutton C, Jackson J, Bennaceur K, Xu Q, Makimanejavali AR, Del Bello F, Piergentili A, Newell DR, Hardcastle IR, Griffin RJ, Lunec J. TP53 mutant MDM2-amplified cell lines selected for resistance to MDM2-p53 binding antagonists retain sensitivity to ionizing radiation. Oncotarget. 2016;7:46203– 46218.

40. Jung J, Lee JS, Dickson MA, Schwartz GK, Le Cesne A, Varga A, Bahleda R, Wagner AJ, Choy E, de Jonge MJ, Light M, Rowley S, Macé S, Watters J. TP53 mutations emerge with HDM2 inhibitor SAR405838 treatment in de-differentiated liposarcoma. Nat Commun. 2016;7:12609.

41. Hata AN, Rowley S, Archibald HL, Gomez-Caraballo M, Siddiqui FM, Ji F, Jung J, Light M, Lee JS, Debussche L, Sidhu S, Sadreyev RI, Watters J, Engelman JA. Synergistic activity and heterogeneous acquired resistance of combined MDM2 and MEK inhibition in KRAS mutant cancers. Oncogene. 2017;36:6581–91.

42. Vassilev LT, Vu BT, Graves B, Carvajal D, Podlaski F, Filipovic Z, Kong N, Kammlott U, Lukacs C, Klein C, Fotouhi N, Liu EA. In vivo activation of the p53 pathway by small-molecule antagonists of MDM2. Science. 2004;303:844–8.

43. Michaelis M, Wass MN, Cinatl J jr. The Resistant Cancer Cell Line (RCCL) Collection. https://research.kent.ac.uk/ibc/the-resistant-cancer-cell-line-rccl- collection/. Accessed 17 Aug 2018.

44. Michaelis M, Agha B, Rothweiler F, Löschmann N, Voges Y, Mittelbronn M, Starzetz T, Harter PN, Abhari BA, Fulda S, Westermann F, Riecken K, Spek S, Langer K, Wiese M, Dirks WG, Zehner R, Cinatl J, Wass MN, Cinatl J Jr. Identification of flubendazole as potential anti-neuroblastoma compound in a large cell line screen. Sci Rep. 2015;5:8202.

45. Weber K, Bartsch U, Stocking C, Fehse B. A multicolor panel of novel lentiviral “gene ontology” (LeGO) vectors for functional gene analysis. Mol Ther. 2008;16:698– 706.

46. Riecken K. Lentiviral Gene Ontology Vectors. http://www.lentigo-vectors.de. Accessed 17 Aug 2018.

47. Voges Y, Michaelis M, Rothweiler F, Schaller T, Schneider C, Politt K, Mernberger M, Nist A, Stiewe T, Wass MN, Rödel F, Cinatl J. Effects of YM155 on survivin levels and viability in neuroblastoma cells with acquired drug resistance. Cell Death Dis. 2016;7:e2410.

48. Li H, Handsaker B, Wysoker A, Fennell T, Ruan J, Homer N, Marth G, Abecasis G, Durbin R; 1000 Genome Project Data Processing Subgroup. The Sequence Alignment/Map format and SAMtools. Bioinformatics. 2009;25:2078–9.

49. Koboldt DC, Zhang Q, Larson DE, Shen D, McLellan MD, Lin L, Miller CA, Mardis ER, Ding L, Wilson RK. VarScan 2: somatic mutation and copy number alteration discovery in cancer by exome sequencing. Genome Res. 2012;22:568–76.

50. Zondlo SC, Lee AE, Zondlo NJ. Determinants of specificity of MDM2 for the activation domains of p53 and p65: proline27 disrupts the MDM2-binding motif of p53. Biochemistry. 2006;45:11945–57.

51. Stiewe T, Haran TE. How mutations shape p53 interactions with the genome to promote tumorigenesis and drug resistance. Drug Resist Updat. 2018;38:27–43.

52. Bouaoun L, Sonkin D, Ardin M, Hollstein M, Byrnes G, Zavadil J, Olivier M. TP53 Variations in Human Cancers: New Lessons from the IARC TP53 Database and Genomics Data. Hum Mutat. 2016;37:865–76.

53. The IARC TP53 Database. http://p53.iarc.fr. Accessed 1 Aug 2018.

54. Prokocimer M, Molchadsky A, Rotter V. Dysfunctional diversity of p53 proteins in adult acute myeloid leukemia: projections on diagnostic workup and therapy. Blood. 2017;130:699–712.

55. Wong TN, Ramsingh G, Young AL, Miller CA, Touma W, Welch JS, Lamprecht TL, Shen D, Hundal J, Fulton RS, Heath S, Baty JD, Klco JM, Ding L, Mardis ER, Westervelt P, DiPersio JF, Walter MJ, Graubert TA, Ley TJ, Druley T, Link DC, Wilson RK. Role of TP53 mutations in the origin and evolution of therapy-related acute myeloid leukaemia. Nature. 2015;518():552–5.

56. Wong TN, Miller CA, Jotte MRM, Bagegni N, Baty JD, Schmidt AP, Cashen AF, Duncavage EJ, Helton NM, Fiala M, Fulton RS, Heath SE, Janke M, Luber K, Westervelt P, Vij R, DiPersio JF, Welch JS, Graubert TA, Walter MJ, Ley TJ, Link DC. Cellular stressors contribute to the expansion of hematopoietic clones of varying leukemic potential. Nat Commun. 2018;9:455.

57. Pommier Y, Leo E, Zhang H, Marchand C. DNA topoisomerases and their poisoning by anticancer and antibacterial drugs. Chem Biol. 2010;17:421–33.

58. Binenbaum Y, Na’ara S, Gil Z. Gemcitabine resistance in pancreatic ductal adenocarcinoma. Drug Resist Updat. 2015;23:55–68.

59. Tamamyan G, Kadia T, Ravandi F, Borthakur G, Cortes J, Jabbour E, Daver N, Ohanian M, Kantarjian H, Konopleva M. Frontline treatment of acute myeloid leukemia in adults. Crit Rev Oncol Hematol. 2017;110:20–34.

